# Optimizing *in utero* electroporation of the developing mouse pons to model diffuse midline glioma

**DOI:** 10.1101/2025.04.06.647423

**Authors:** Madisen S. Mason, Hosbaldo Morales Murillo, Madisyn G. Dudek, Samantha E. Maher, Santos J. Franco

## Abstract

The leading cause of brain cancer-related death in children is diffuse midline glioma (DMG). A particularly aggressive DMG subtype is pediatric diffuse intrinsic pontine glioma (DIPG), which is caused by the histone mutation H3.3K27M. Because of its diffuse growth and location in a critical brainstem structure, therapeutic options are limited and DIPG is considered universally fatal. Lack of appropriate animal models has hindered our understanding of the developmental origins and progression of DIPG, which in turn has limited development of effective therapeutics. To address this barrier, we optimized an *in utero* electroporation method to model DIPG *in vivo* in the developing mouse pons. In this protocol, we use *in utero* electroporation to express canonical DIPG mutant oncogenes in neural progenitors lining the 4^th^ ventricle, which give rise to cells in the pons. As the embryos continue to develop *in utero* and then postnatally, they develop large diffuse brainstem tumors with molecular characteristics of pediatric DIPG, allowing us to model the formation and progression of this deadly pediatric brain cancer.

## INTRODUCTION

Tumors of the central nervous system are the most common cancer-related cause of death in children. Pediatric high-grade gliomas are particularly aggressive and deadly, accounting for nearly half of all deaths caused by pediatric brain tumors [1]. Importantly, high-grade gliomas in children differ significantly from those that develop in adulthood in terms of their cellular origins and biological drivers [2,3]. These differences are reflected in the fact that clinical trials and treatments based on adult high-grade gliomas have shown no significant improvement in clinical outcomes in children [4]. Tumors also arise in a specific spatio-temporal pattern, typically in middle childhood [5], suggesting that tumor cells arise from dysregulation of a normal developmental process. Because of its location, current treatment options are limited, and prognosis is poor [6].

In recent years, significant advancements in patient-derived cell lines and animal models have provided a solid foundation to expand our understanding of the molecular underpinnings of these tumors. Gain-of-function mutations in genes encoding histone 3 (H3) variants H3.1 and H3.3 are found in a substantial proportion of pediatric high-grade gliomas, and these mutations are always present in diffuse midline glioma (DMG, H3K27M-altered) [7]. Mutation of lysine 27 to methionine on H3 (H3K27M mutations) causes loss of trimethylation at this position, resulting in aberrant epigenetic regulation in affected cells [8,9]. The resulting transcriptional changes can significantly impact neural progenitor cells that rely on precise gene regulatory networks to produce the correct types of cells in proper numbers. H3K27M mutations in these tumors are often combined with genetic loss of *TP53* and amplification of *PDGFRA* [2,3,10–12]. Together, these studies indicate that the primary driver of DMG is epigenetic and transcriptional dysregulation in neural progenitors and glial precursors, combined with suppression of apoptosis and activation of growth factor pathways.

Most current animal models of DIPG involve orthotopic xenografts of human tumor cells into adult immunodeficient mice. Though these models produce tumors, they require immunodeficient mice to generate tumors, and the xenografted tumors grow in adult mice, not in the developing brain as in human patients. A few genetically engineered mouse models of DMG have been developed to generate tumors that form in early embryonic and postnatal development in animals with competent immune systems [13,14]. However, studies using genetically engineered mice have been sparse, probably because of the challenges of engineering multiple inducible mutant alleles into the same mouse line. Recently, *in utero* electroporation (IUE) has been used to generate spatiotemporally defined gliomas with specific genetic drivers for pediatric DMG in an immunocompetent setting [15,16]. This strategy overcomes many of the limitations of prior approaches but seemed to face some technical hurdles that made the IUE approach challenging. Indeed, only one follow-up study has been published since these initial reports [17]. IUE has proven to be an invaluable tool for the study of brain development, and our group has utilized this approach extensively [18–21]. Here we describe modification and optimization of an *in utero* electroporation protocol to study the developmental origins of DIPG. Previous protocols have already described the materials and general steps of IUE of the cerebral cortex [22,23], so we do not readdress those details here. This protocol describes an adaptation to target the developing pons using a third electrode in addition to standard forceps-type electrodes. By adjusting the position and polarity of the electrodes, neural progenitors in the embryonic pons primordium can be electroporated, resulting in spatially specific tumor formation in the mouse developing pons.

## MATERIALS AND METHODS

The protocol described in this peer-reviewed article is published with full details on protocols.io [24], (**dx.doi.org/10.17504/protocols.io.5jyl8dy7dg2w/v1**) and is included for printing as (S1 File) with this article.

### Expression Plasmids

PB-pCIG has been previously described [21]. TRIONCO plasmid was custom synthesized and purchased from GenScript Biotech. The plasmid comprises mouse *H3f3a* with a K27M mutation fused to GFP at the C-terminus (H3^K27M^-GFP), a P2A sequence, mouse *Trp53* with a dominant-negative R270H mutation fused to a V5 tag at the C-terminus (p53^R270H^-V5), a T2A sequence, and mouse *Pdgfra* with a constitutively-active D842V mutation (PDGFRA^D842V^), all cloned into the PB-pCIG backbone missing the IRES-GFP. The plasmid was confirmed by whole-plasmid DNA sequencing (plasmidsaurus). Full sequence available upon request. PB-DoubleUP was cloned by inserting a PCR fragment from Double UP sfGFP to mScarlet [25] into the PB-pCIG backbone missing the IRES-GFP using NEBuilder HiFi (New England Biolabs). CMV-mPB plasmid [18] expressing *piggyBac* transposase was co-electroporated to permit stable integration of the plasmids into electroporated progenitors.

### Immunohistochemistry and Confocal Imaging

Embryonic brains were fixed in 4% paraformaldehyde (PFA) overnight at 4°C. Brains were sectioned obliquely (see Fig 1F) at 100 µm with a vibrating microtome. Free-floating sections were placed in 24-well plates and blocked with 500 μl of 10% donkey serum and 0.2% Triton X-100 in 1× PBS for 2 h at RT. Blocking solution was then removed, and sections were incubated with primary antibodies in 10% donkey serum and 0.2% Triton X-100 in 1× PBS overnight (16 h) at 4°C. Primary antibody solution was then removed, and sections were washed at RT with 1× PBS three times for 5 min each. After washing, sections were incubated with secondary antibodies in 1× PBS for 1 h at RT. Sections were then washed again using 1× PBS three times for 5 min each. Sections were mounted on glass slides with ProLong Diamond Antifade Mountant (Thermo Fisher Scientific). Images were captured using a Zeiss LSM 900 laser scanning confocal microscope in Airyscan 2 Multiplex 4Y mode at 20× or 40× magnification.

**Fig 1.**
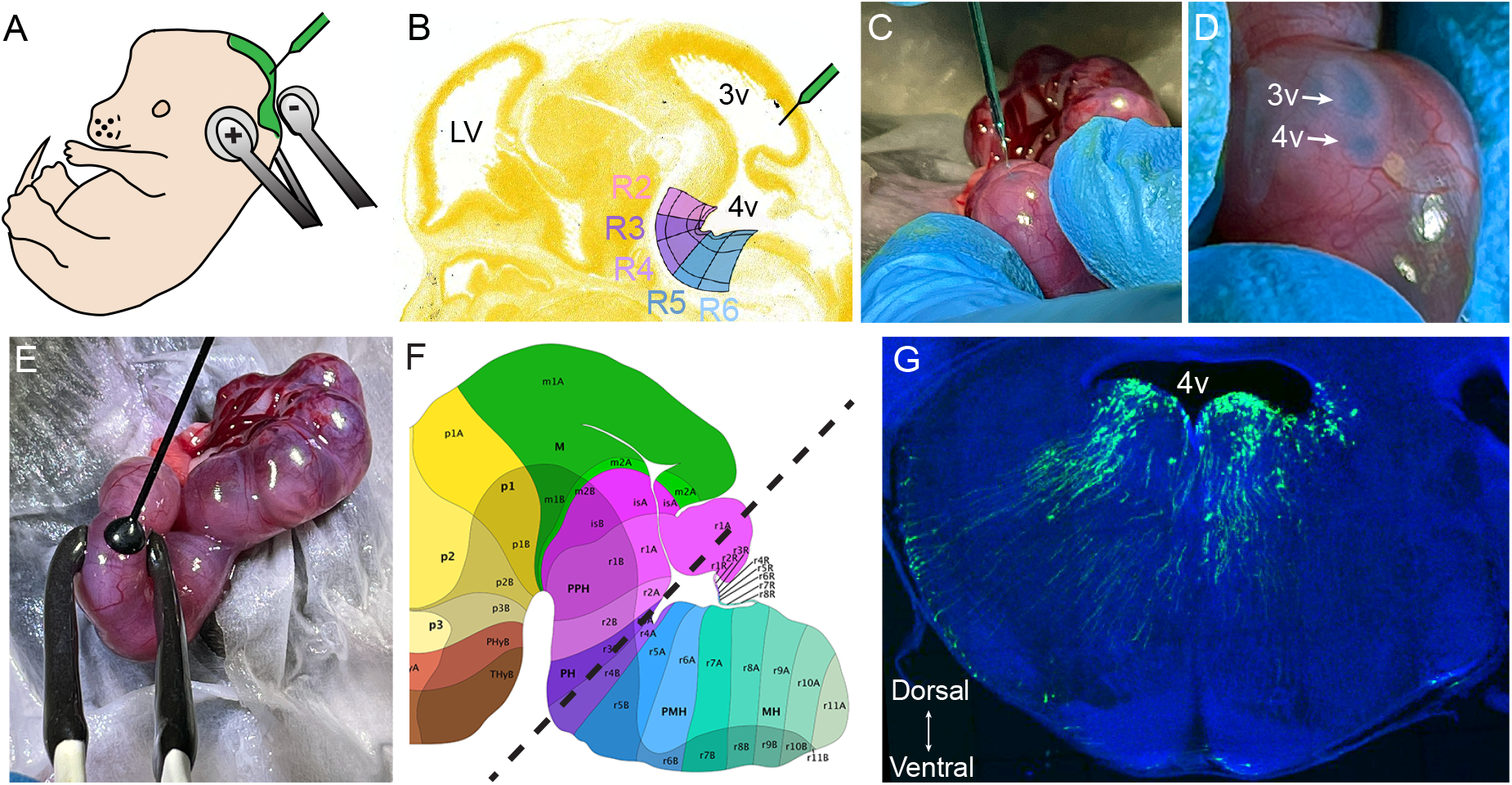
Targeting the embryonic pons by in utero electroporation using triple electrodes. **(A)** Schematic of IUE with injection into the 4th ventricle and triple electrode placement. **(B)** Feulgen-HP yellow stain of a sagittal section of a mouse brain at E13.5, overlaid with anatomical annotations for target rhombomeres 2-6 (R2-6). From the Allen Reference Atlas – Developing Mouse Brain [brain atlas], available from atlas.brain-map.org. **(C-E)** Images of an IUE procedure showing injection of DNA mix into an embryo using a glass microcapillary needle (C), filling of the 3rd and 4th ventricles (arrows) with DNA mix, visualized by Fast Green dye (D), and placement of triple electrodes around the injection sites (E). **(F)** Atlas of a mouse brain at E17.5, showing anatomical annotations from the Allen Reference Atlas – Developing Mouse Brain [brain atlas], available from atlas.brain-map.org. Dashed line shows oblique angle of sectioning for image in (G). **(G)** Results of an IUE of control sfGFP at E15.5, dissected at E17.5 and cut obliquely as in (F). Image shows electroporated neural progenitors with cell bodies in the ventricular zone of the 4th ventricle and their radial processes spanning the entire pons. Abbreviations: LV, lateral ventricles; 3v, 3rd ventricle; 4v, 4th ventricle.

Antibodies used for immunostaining are listed in **Biological and Chemical Reagents** table below. The concentration of each primary antibody used was the following: mouse anti-H3K27M (1:500), mouse anti-H3K27me3 (1:500), goat anti-OLIG2 (1:1000), rabbit anti-SOX2 (1:500), rat anti-PDGFRA (1:1000), chicken anti-GFP (1:500), and DAPI (1:10,000). Donkey secondary antibodies conjugated to Alexa Fluor 488, Rhodamine Red-X, or Alexa Fluor 647 were used at 1:500.

### Biological and Chemical Reagents

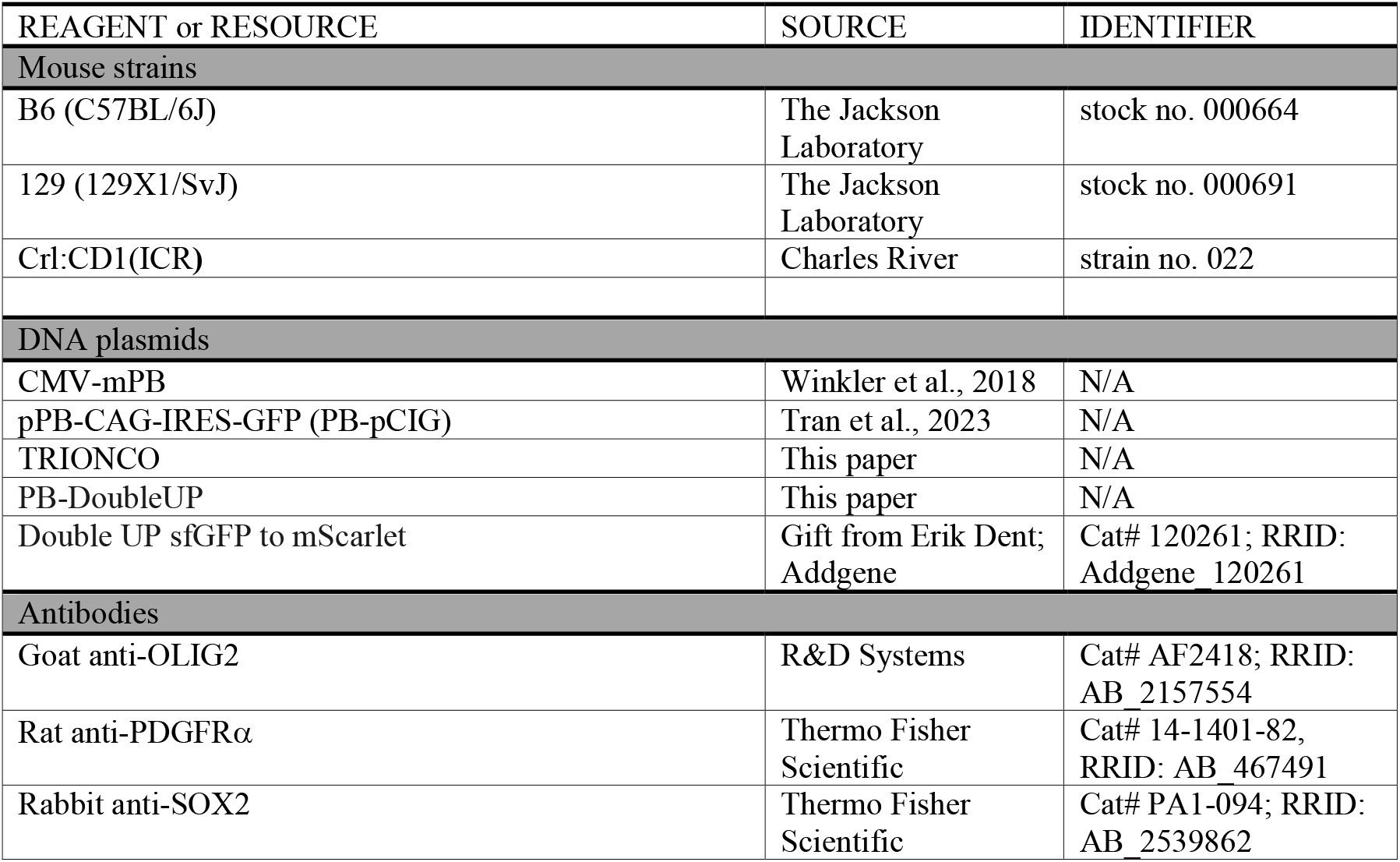
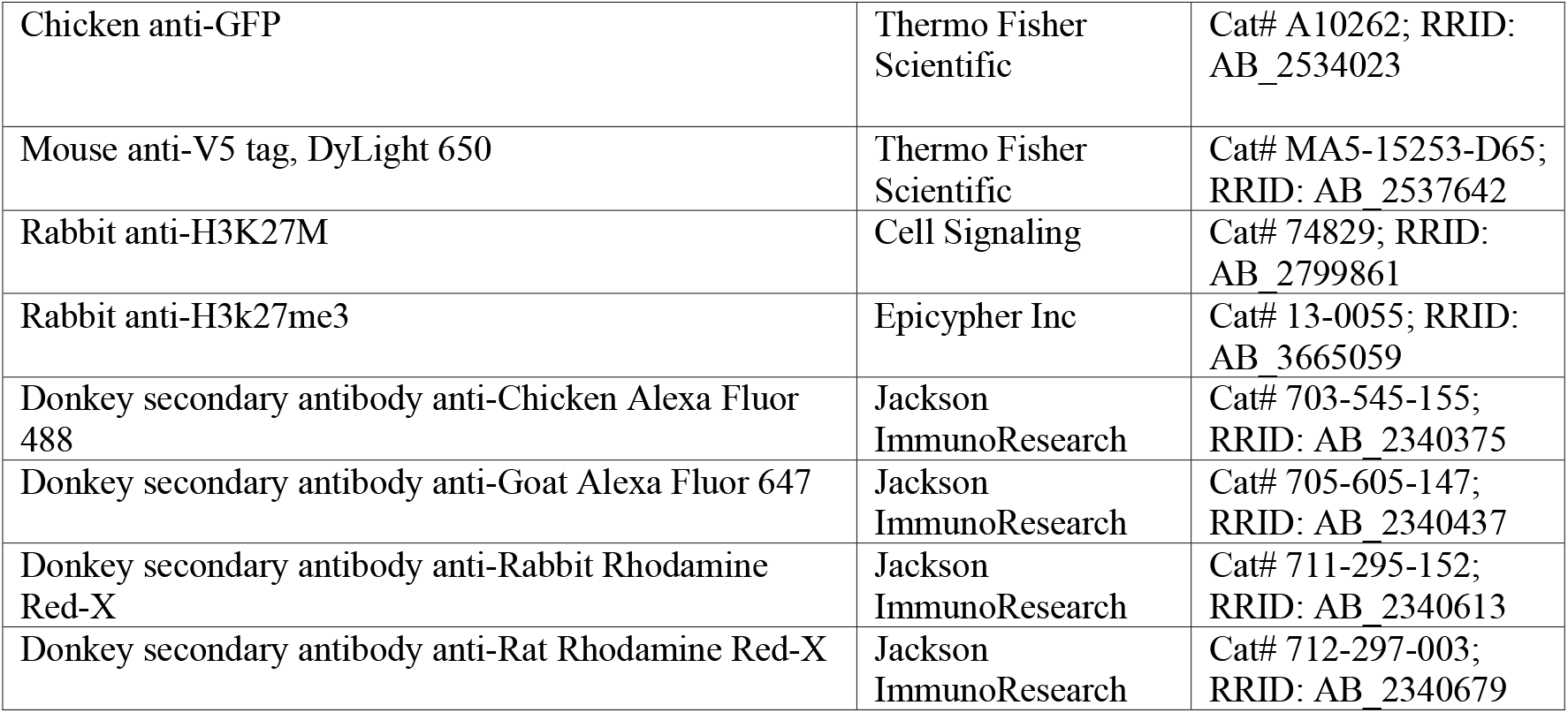

## RESULTS AND DISCUSSION

### Triple Electrode Approach to Electroporate 4^th^ Ventricle Progenitors *In Utero*

Using the 4^th^ ventricle *in utero* electroporation protocol we described in the material and methods, we injected an electroporation mix containing a super folder GFP-expressing control vector (PB-DoubleUP) and *piggyBac* transposase (CMV-mPB) into the caudal end of the 3^rd^ ventricle (**Fig 1A and Fig 1C**) of embryos at embryonic day 15.5 (E15.5). Angling the injection needle caudally toward the 4^th^ ventricle allowed the plasmid mix to flow into the 4^th^ ventricle via the cerebral aqueduct (**Fig 1A and Fig 1B**). Proper injection into the 4^th^ ventricle was visualized by the Fast Green dye in the electroporation mix and identified as a small diamond-shaped spot caudal to the 3^rd^ ventricle, which was also often filled (**Fig 1C**). The ventricular zone along the 4th ventricle contains neural progenitors in rhombomeres 2-4 that give rise to the developing pons (**Fig 1B**). Using a triple-electrode setup [26], we electroporated the pons ventricular zone progenitors. We placed tweezer-style cathode electrodes on the lateral sides of the embryo’s upper neck on either side of and just ventral to the 4^th^ ventricle, and we placed the single anode electrode directly on the top of the 4^th^ ventricle (**Fig 1E**). This strategy allowed us to drive the electric current from the electroporator from dorsomedial to ventrolateral, thus introducing the plasmid DNA into progenitors in the ventricular zone of the 4^th^ ventricle. We dissected brains at E17.5 and cut oblique sections through the developing pons (**Fig 1F**). Confocal imaging revealed GFP-expressing neural progenitors in both the basal and alar plates of rhombomeres 2-4, with their cell bodies located in the ventricular zone and long radial processes spanning to the ventral surface of the pons (**Fig 1G**). Thus, this protocol allows efficient electroporation of neural progenitors throughout the ventricular zone of rhombomeres 2-4 that give rise to neurons, astrocytes, oligodendrocytes, and ependymal cells of the pons tegmentum [27].

Use of the triple electrode approach allowed us to optimize targeting of neural progenitor cells lining the pons ventricular zone. This strategy can also electroporate progenitors in the lower rhombic lip if the tweezer electrodes are placed slightly more dorsally around the 4^th^ ventricle (data not shown). The rhombic lip is a germinal matrix lining the dorsolateral edges of the 4^th^ ventricle, near the roofplate, that produces a highly diverse populations of cells in the brainstem and cerebellum [28]. The rhombic lip is separated into two distinct regions: the upper (rostral or cerebellar) rhombic lip in rhombomeres 1 and 2 that give rise to cells in the cerebellum [29], and the lower (caudal or hindbrain) rhombic lip in rhombomeres 2-7 that give rise to neurons that migrate tangentially to populate precerebellar nuclei, including pontine nuclei in the basilar pons [30]. Changing electrode placement or polarity can more effectively target the upper and/or lower rhombic lip if desired. We have also found that using just the tweezer-style electrodes alone, with one side as the cathode and the other the anode, can target either the left or right side of the rhombic lip depending on electrode placement.

### Validation of the TRIONCO Plasmid

After demonstrating that we could target the pons ventricular zone, we next wanted to use this technique to generate DMG tumors in the pons to model DIPG. Toward this goal, we first generated a plasmid containing three canonical DIPG oncogenes: 1) H3.3^K27M^, a point mutation that globally reduces histone methylation on lysine 27 (H3K27me3), 2) p53^R270H^, a dominant-negative version of p53 that blocks apoptosis, and 3) PDGFRA^D842V^, a constitutively-active version of platelet-derived growth factor receptor alpha (**Fig 2A and Fig 2B**). This combination of oncogenes has been shown previously to cause high-grade gliomas in a mouse *in utero* electroporation model [16]. However, in prior studies the oncogenes were encoded on separate plasmids and co-electroporated. Simultaneously electroporating multiple plasmids at similar molar ratios can result in relatively high co-expression rate [31,32]. However, we found in early pilot experiments using the co-electroporation approach that many cells were expressing only the constitutively active PDGFRA^D842V^, and that this could result in significant hyperplasia even in the absence of H3.3^K27M^ or p53^R270H^ (data not shown). Therefore, we encoded all 3 oncogenes on a single plasmid, referred to here as TRIONCO, to ensure co-expression in electroporated cells.

**Fig 2.**
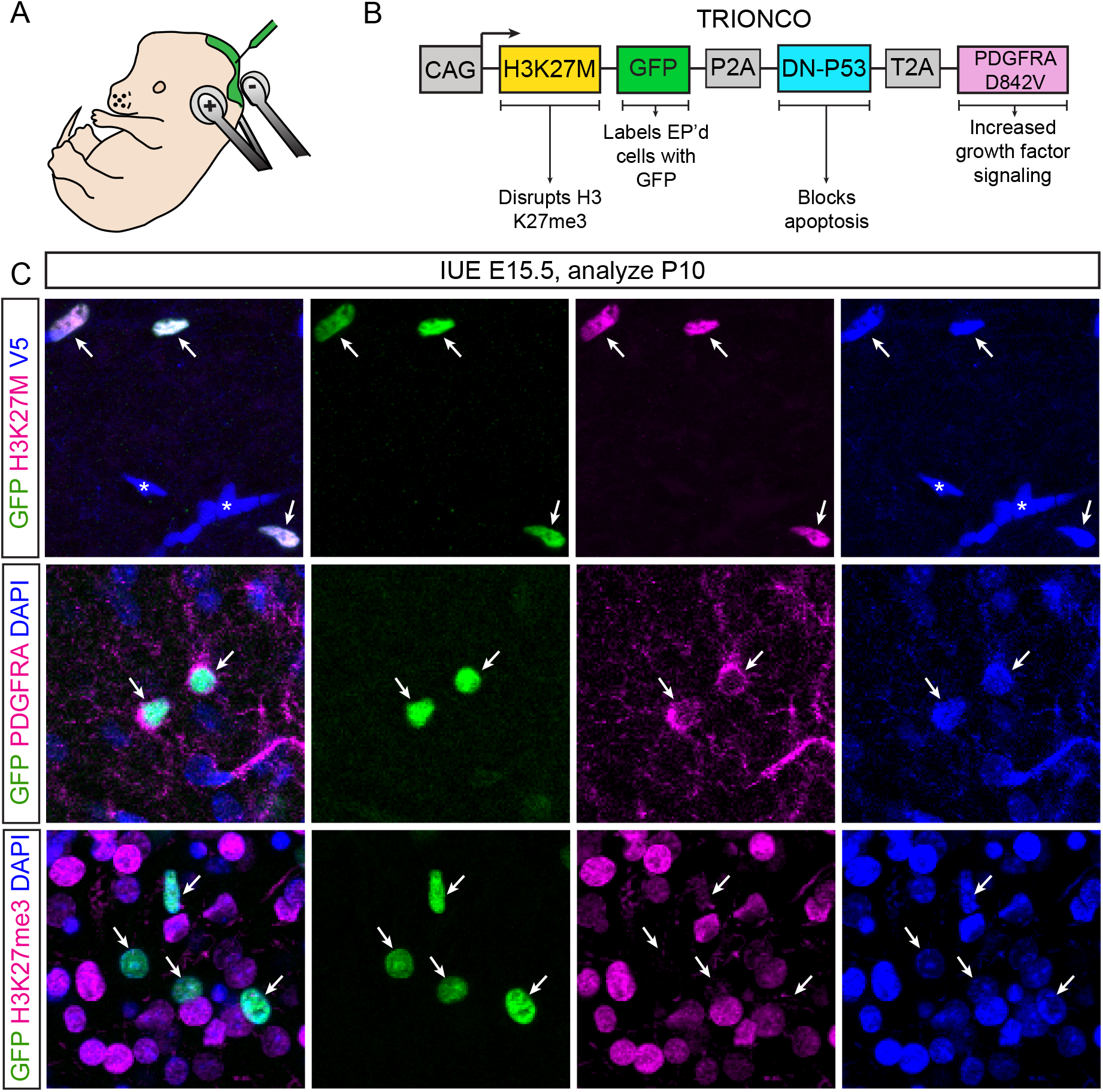
Validation of oncoprotein expression in brains electroporated with the TRIONCO plasmid. **(A)** Schematic of IUE with injection into the 4th ventricle and triple electrode placement. **(B)** Schematic of TRIONCO plasmid to express DMG driver oncogenes, with descriptions of the functions of the encoded cDNAs. **(C)** Representative images of brains electroporated at E15.5 and analyzed at P10. Oblique sections were cut and stained for GFP (green), H3K27M (magenta), and V5 (blue) [top row], GFP (green), PDGFRA (magenta), and DAPI (blue) [middle row], or GFP (green), H3K27me3 (magenta), and DAPI (blue) [bottom row]. Merged channels shown in the left column, individual images at right. Arrows point at GFP+ cells that also co-express the other markers.

To validate our plasmid, we first electroporated TRIONCO at E15.5 and allowed the embryos to develop to postnatal day 10 (P10). We then dissected the electroporated brains and performed immunohistochemistry to verify expression of the encoded oncogenic proteins. Staining for antibodies against the mutant H3.3^K27M^ protein, the V5 tag fused to p53^R270H^, and PDGFRA demonstrated co-expression of all three proteins in GFP+ electroporated cells (**Fig 2C**). We also verified lower levels of H3K27me3 signal in GFP+ cells compared to neighboring cells that were not electroporated (**Fig 2C**), indicating that the H3K27M mutation was able to reduce H3K27me3 levels. Together, these data demonstrate proper expression of all 3 oncogenes after TRIONCO electroporation, leading to reduced histone 3 lysine 27 trimethylation levels.

### Generation of Diffuse Midline Glioma Tumors

We next analyzed electroporated brains at different timepoints after TRIONCO electroporation at E15.5 to assess cellular expansion and tumor growth. Two days after electroporation, at E17.5, a small number of GFP+ electroporated cells were found bilaterally in and near the ventricular zone of the 4^th^ ventricle (**Fig 3A**). The numbers of GFP+ cells were similar in TRIONCO (**Fig 3A**) and control PB-DoubleUP (**Fig 1G**), indicating relatively low levels of proliferation of electroporated cells at this timepoint. By P10, that small number of GFP+ cells has increased to a several hundred cells that were distributed diffusely away from the ventricle, indicating substantial proliferation and migration into the developing pons. By P60 there was a substantial expansion of GFP+ electroporated cells numbering in the thousands (**Fig 3C**). The GFP+ cells were located diffusely throughout the dorsal-ventral and medial-lateral axes of the mature pons, with a higher concentration along the midline (**Fig 3C**).

**Fig 3.**
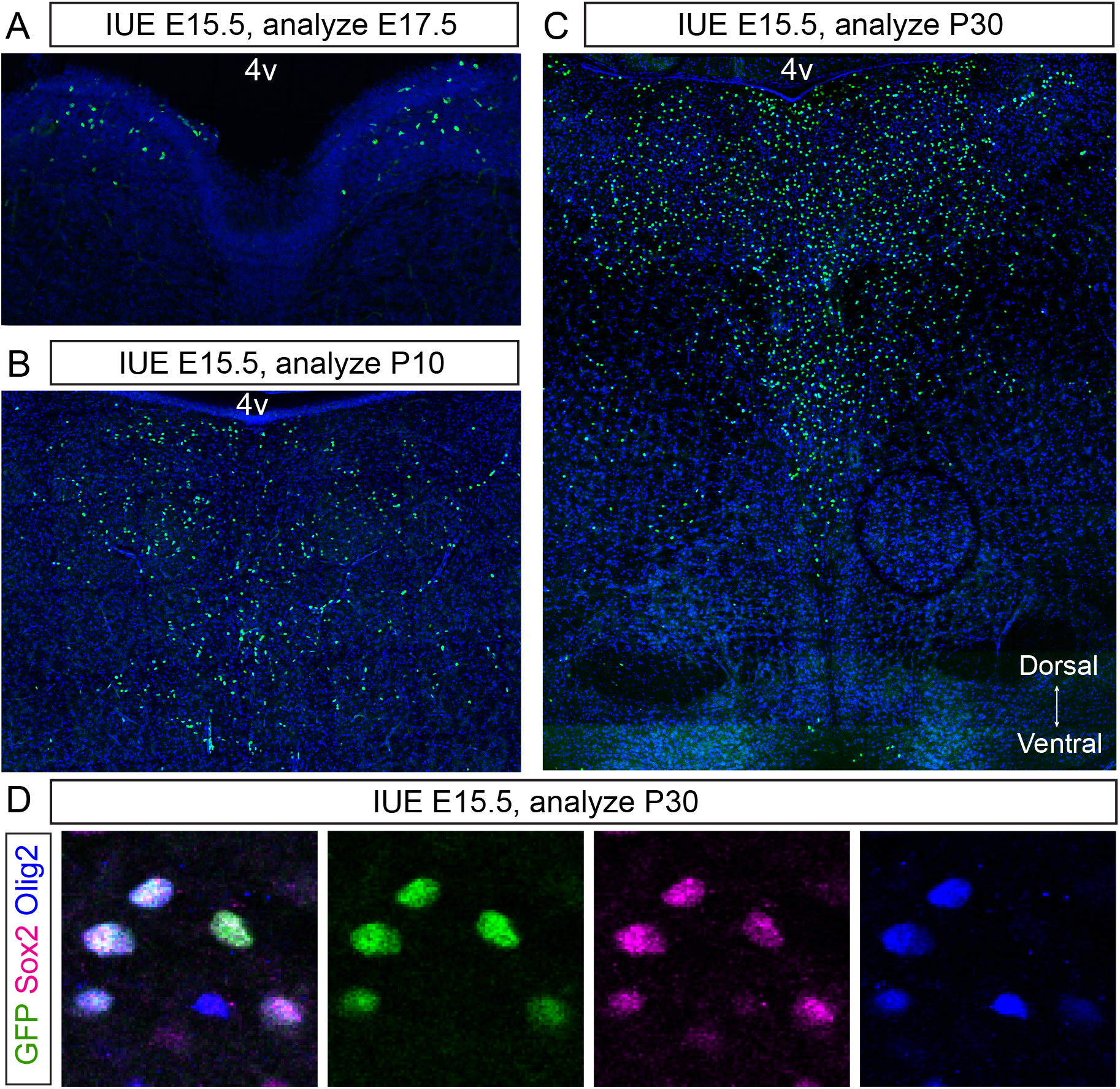
Mice electroporated with TRIONCO progressively develop diffuse tumors with molecular characteristics of DIPG. **(A-C)** Representative images of brains electroporated with TRIONCO at E15.5 and analyzed at various ages after IUE. At E17.5 (A) there are dozens of electroporated cells (green nuclei) per section. By P10 (B) electroporated cells have expanded into hundreds per section. At or P30 there are thousands of tumor cells per section, diffusely scattered throughout the pons. **(D)** Representative images of brains electroporated at E15.5 and analyzed at P30, showing co-expression of DIPG markers in GFP+ electroporated cells. Oblique sections were cut and stained for GFP (green), SOX2 (magenta), and OLIG2 (blue). Merged channels shown in the left column, individual channels at right.

Immunohistochemistry for canonical DMG markers SOX2 and OLIG2 showed that nearly all GFP+ cells electroporated with TRIONCO were SOX2+, with the majority of those also expressing OLIG2 (Figure 3D). Thus, cells electroporated with TRIONCO using the triple electrode protocol expand diffusely throughout the pons and express canonical DMG markers found in human tumors. Implementation of the optimized protocol presented here will help facilitate use of the pons *in utero* electroporation model for studying aspects of diffuse midline glioma formation and growth that are not possible using other currently available approaches.

## Additional Considerations

We have successfully used this protocol to electroporate embryos between E12.5 and E15.5. Injection of the electroporation mix and placement of electrodes can be more challenging at E12.5-E13.5 compared to E14.5-E15.5, leading to lower rates of successfully electroporated brains, especially for less experienced surgeons. Birth of electroporated pups typically occurs on gestational day 19.0-19.5, so it is recommended to minimize disturbing the pregnant dam for about 1 week after surgery to facilitate best survival rates. In a typical experiment >80% of injected embryos will be born and nearly all surviving pups will develop DIPG tumors, although reproducibility of this protocol is highly dependent on experience and skill levels.

We have had success performing this protocol on both the CD1 outbred strain and a hybrid of 129;B6 inbred strains, each of which has some advantages and disadvantages. Females from both strains tend to be good mothers, which facilitates pup survival. CD1 mice have larger average litter sizes, which can increase numbers of electroporated mice but also increases variability, surgery time, and chances of complications. 129;B6 hybrids have smaller litter sizes than CD1, but more consistent developmental stages across all embryos in the litter. For both strains, including enrichment and enhanced diets can positively impact pup survival. We have not attempted this specific protocol on a pure C57BL6/J inbred strain, although our prior experience with electroporation of the lateral ventricles indicated that females of this strain do not care for their pups very well after *in utero* electroporation surgery and therefore survival of electroporated pups is extremely low.

To date we have not seen obvious differences in tumors formed in mice electroporated at E13.5 compared to E15.5. However, neural progenitors give rise to different subtypes of cells at different developmental timepoints, raising the possibility that timing of electroporation can impact cell of origin and therefore tumor characteristics. Thus, electroporation timing should be determined for each experimental paradigm depending on study goals. Likewise, the time of analysis will be highly study dependent. We have found that cell numbers continue to expand in TRIONCO tumors at least through P60, although we have not yet carefully analyzed tumor growth over time in a quantitative manner. In future studies it will be useful to incorporate a strategy to co-electroporate an additional plasmid encoding Luciferase, allowing tumor growth to be tracked longitudinally by *in vivo* bioluminescence imaging.

## Ethics declarations

Experiments involving animals were performed according to the guidelines from the Institutional Animal Care and Use Committee of the University of Colorado–Anschutz Medical Campus and were approved in CU-AMC IACUC protocol #19.

## Supporting Information

S1 File: Step-by-step protocol, also available on protocols.io.

dx.doi.org/10.17504/protocols.io.5jyl8dy7dg2w/v1

## Acknowledgements

This work was supported by the Cancer League of Colorado (S.J.F.), the Colorado Child Health Research Institute Marilyn Hodges Wilmerding Research Pilot Award (S.J.F.), and the Summer Research Training Program (H.M.M.). We thank the following people for helpful scientific discussions: Dr. Adam Green, Dr. Siddhartha Mitra, and Salvador Guerra.

## Authors’ Contributions

Conceptualization, M.S.M., S.J.F.; Investigation, M.S.M., H.M.M., M.D., S.M., S.J.F.; Writing – Original Draft, M.S.M., S.J.F.; Visualization, M.S.M., S.J.F.; Funding Acquisition, S.J.F.

