## Supplemental File S1 for "Optimizing *in utero* electroporation of the developing mouse pons to model diffuse midline glioma"

Mar 03, 2025

### *In utero* electroporation of mouse pons via the 4th ventricle

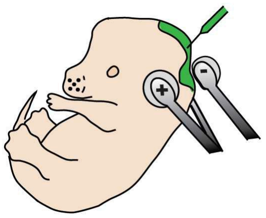

DOI
**dx.doi.org/10.17504/protocols.io.5jyl8dy7dg2w/v1**

Madisen S Mason[1], Santos J Franco[1]

[1]University of Colorado Anschutz Medical Campus

Madisen S Mason
University of Colorado

OPEN ACCESS

**DOI:** dx.doi.org/10.17504/protocols.io.5jyl8dy7dg2w/v1

**Protocol Citation:** Madisen S Mason, Santos J Franco 2025. In utero electroporation of mouse pons via the 4th ventricle. **protocols.io** https://dx.doi.org/10.17504/protocols.io.5jyl8dy7dg2w/v1

**License:** This is an open access protocol distributed under the terms of the **Creative Commons Attribution License**, which permits unrestricted use, distribution, and reproduction in any medium, provided the original author and source are credited

**Protocol status:** Working
**We use this protocol and it's working**

**Created:** February 20, 2025

**Last Modified:** March 03, 2025

**Protocol Integer ID:** 123176

**Keywords:** in utero electroporation, pons, 4th ventricle, mouse

**Funders Acknowledgements:**
**Cancer League of Colorado**
**Grant ID: 220552-SF**
**Colorado Child Health Research Institute**
**Grant ID: R9010L**

#### Abstract

*In utero* electroporation of the developing nervous system is a powerful approach to studying neural development *in vivo*. Most studies using *in utero* electroporation have focused on electroporating neural progenitors lining the lateral ventricles, which give rise to cells in the forebrain. Here, we provide a step-by-step protocol to perform *in utero* electroporation of neural progenitors lining the mouse 4th ventricle, which give rise to cells in the pons. This method can be used to study normal development of the pons and to model human diseases like pediatric high-grade gliomas.

### Materials

**Mice**

E13.5 pregnant females, 129;B6 hybrid strain (F1 offspring of 129X1/SvJ females x C57BL6/J males)

- C57BL6/J, The Jackson Laboratory stock no. 000664
- 129X1/SvJ, The Jackson Laboratory stock no. 000691

**Equipment and Tools**

- Micropipette puller, Sutter model P-97
- EZ-Anesthesia Microflex Small Animal Anesthesia System w Heat, BrainTree Scientific
- Electroporator BTX ECM 830, with foot pedal and cables
- Y-Cable, 2 socket, 2 mm Female to 1 socket, 2 mm Male (monopole) plug, Bulldog Bio C118
- Tweezertrode-style electrodes, 3 mm, platinum, Bulldog Bio CUY650P3
- Single disk electrode, 3 mm, platinum, Bulldog Bio CUY700P3L
- Hot bead sterilizer, 250°C, Fine Science Tools 18000-45
- Heating block (dry bath) with 50mL conical tubes adapter
- Surgical lamp with flexible arms (Halogen 20W or LED equivalent)
- Cordless Hair clippers
- Graefe Forceps - 10cm straight, 0.8mm tip, Fine Science Tools 11050-10
- Ring Forceps, 9cm, ID-2.2mm, OD-3mm, Fine Science Tools 11103-09
- Trident II Spring Scissors, 8cm straight, cutting edge 8mm, tip 0.2mm, Fine Science Tools 15110-08
- Olsen-Hegar Needle Holders w Suture Cutters, Fine Science Tools 12002-12
- Instrument sterilization case with spiked silicon mat, autoclaveable, Fine Science Tools 20810-02

**Reagents**

- Plasmid DNA, endotoxin-free maxi prep, highly concentrated (> 2 mg/ml)
- Fast Green FCF (0.1% in endotoxin-free, nuclease-free water), ThermoFisher A16520-06
- Endotoxin-free, nuclease-free water or Tris-EDTA
- Borosilicate Glass capillaries O.D. 1mm, I.D. 0.58m, 6cm length, World Precision Instruments #1B100-6
- Isoflurane, 99.9% USP, Piramal Critical Care #6679401725
- Povidone-iodine (Betadine) prep wipe, 10% USP, ThermoFisher 666970
- Isopropanol prep wipe, 70% USP, ThermoFisher 06-669-62
- Paralube ocular lubricant, Ferring Laboratories #00574402535
- HBSS (1x), ThermoFisher 14175095
- 50 mL conical tube for HBSS
- Transfer pipet, 3 mL, sterile, individually wrapped
- Meloxicam, Selleck Chemical 71125-38-7
- 1.5 mL conical tube for Meloxicam
- Syringe, 1 mL
- Needle, 26G x 3/8" with intradermal bevel
- Sutures, Plain Gut, 6-0, P-1, 18" (774PG) or Prolene, 6-0, P-1, 18" (8697G)
- Sterile Latex Powdered Surgical Gloves

#### Before start

Before you start, prepare DNA injection mix, beveled microcapillary needles, analgesic, and surgical tools.
Dilute selected endotoxin-free plasmid DNA in sterile nuclease-free water to 1-2 µg/µl and add Fast Green dye to the DNA mix. Pull and bevel microcapillary needles for plasmid injection - 1 primary needle and 1-2 backup needles per injection mix. Prepare Meloxicam aliquots. Autoclave surgical tools.

Refer to Courchet et al. 2020 for more comprehensive details on preparation of materials and full surgical procedure. This protocol is an adaptation specific for injecting the 4th ventricle and electroporating progenitors for the pons, but closely follows the general surgical details in the published manuscript.

CITATION

Géraldine Meyer-Dilhet, Julien Courchet (2020). In Utero Cortical Electroporation of Plasmids in the Mouse Embryo. STAR Protocols.
LINK
10.1016/j.xpro.2020.100027

#### Prepare the surgical area.

10m

1 If performing multiple consecutive surgeries, turn on bead sterilizer. Use bead sterilizer to sterilize tools between surgeries. When temperature reaches 250°C, sterilize surgical tools (Graefe forceps, ring forceps, spring scissors, and Olsen-Hegar needle holders).

2 Place sterile tools on sterile silicon mat.

3 Turn on water heating circulation pump to heat the surgical bed and cage warming pad (37°C). 10m

4 Place fresh 50 mL conical of HBSS into dry bath block and turn on warmer to low (37°C). Open sterile transfer pipette and place in 50 mL conical of HBSS.

5 Turn on the electroporator. Set electroporator to 35V (E13.5), 40V (E14.5), or 45V (E15.5), 50 ms pulse length, and 4 pulses with 950 ms intervals. Plug Y-cable into the negative (black) port and plug the tweezer electrode male terminals into the Y-cable female plugs. Plug the single red cable into the positive (red) port and plug and the single stick electrode male terminal into the single red cable female plug.

6 Plug the $O_2$ connector into the wall port and set the $O_2$ flow meter on the anesthesia setup to "OFF".

7 Connect the anesthesia machine and fill vaporizer with isoflurane, if necessary. Ensure the correct tube is connected to the induction chamber.

8 Weigh charcoal filter and connect to waste gas scavenger system. Set waste gas scavenger system vacuum settings to 14 (induction chamber line) and 4 (nose cone line).

9 Resuspend Meloxicam aliquot with 190 µL HBSS (2 mg/kg body weight).

10 Set out 2 isopropanol prep wipes and 1 povidone-iodine prep wipe.

#### Anesthetize and prepare the pregnant mouse.

8m

11 Weigh pregnant dam to calculate proper amount of analgesic to administer.

12 Turn on waste gas scavenger system.

13 Induce pregnant dam in induction chamber with oxygen flow meter at 1.05 L/min and isoflurane flow at 5% until the animal is sedated. 2m

13.1 This step will take on average 60-90 seconds depending on the animal's age, weight, and body fat. Carefully monitor to avoid a lethal isoflurane overdose.

14 Once sedated, remove mouse from induction chamber and place it face up (dorsal decubitus) and quickly treat both eyes with Puralube eye lubricant before placing animal's face in nosecone. 30s

14.1 Be sure to switch isoflurane flow from the induction chamber to the nosecone so that oxygen and isoflurane are diverted to nosecone.

15 Tape mouse front paws to surgical bed, gently with science tape. This is to prevent the nose from being pulled out of the nosecone during surgery. 30s

15.1 Do not pull the forelimbs too tight during taping. Stretching the limbs too tightly can impair the mouse's breathing.

16 Slowly decrease Isoflurane to 3.5%. Monitor mouse and frequently check the absence of reflex by pinching the hind paw and tip of the tail to ensure deeply anesthetized state.

17 Assemble needle (30G x 1/2") and syringe (1mL) and pull up appropriate amount of diluted Meloxicam. Final dose 2mg/kg body weight. Subcutaneously inject diluted Meloxicam in mouse under arm. 30s

18 Shave abdomen of the mouse centered around the navel. Prominent anatomical markers are the 3rd and 4th mammary glands. 1m

19 Sterilize the freshly shaven area with alternated cleaning. Start first with an isopropanol swab followed by povidone-iodone. Repeat one final time with an isopropanol swab. 30s

20 Once again, decrease Isoflurane to 2.5% and check the absence of reflex.

21 With your non-dominant hand use Graefe forceps to gently raise the skin above umbilic. With your dominant hand use the spring scissors to make a vertical incision about 1-2 cm in length. You should be able to easily see the white line joining the two abdominal muscles. 30s

22 Again, with Graefe forceps in non-dominant hand, gently raise the abdominal wall and use the spring scissors to make a very small incision (1mm). This will air to enter the abdomen and vital organs to separate from the muscle. Then perform a 1-2 cm long incision along the white line. 30s

23 Place surgical drape across the mouse with opening lining up with incisions made in previous steps. Apply warmed HBSS with transfer pipet into mouse opening to keep the uterine horn damp.

24 Using Graefe forceps in non-dominant hand and ring forceps in dominant hand, gently expose one uterine horn. 2m

24.1 Do not pull out the first embryos you see. Typically these are the embryos closest to the cervix and cannot move much. Instead, target one of the embryos that is a few spots away from the cervix and slowly pull the uterus out one embryo at a time.

24.2 Use the ring forceps to gently grab uterine tissue between embryos, while avoiding blood vessels that line the uterine horns.

24.3 Pull out one side of the uterine horn until you see the ovary. Count the number of embryos and document on the electroporation record sheet. Then repeat on the other side.

24.4 Assess viability of the embryos. Healthy embryos will be light pink and have clear amniotic fluid. Unhealthy embryos can be completely white in color or have cloudy amniotic fluid. Skip unhealthy embryos to reduce cross-contamination.

25 Wet the uterine horns with warmed HBSS.

#### Inject and electroporate the embryos.

10m

26 Fill pulled and beveled microcapillaries with prepareed plasmid mix in the back of the microcapillary via capillary action. Fill needles with around 20µL and insert needle into silicone holder end of aspirator tube. Slowly expel the plasmid mix to the sharp tip of the microcapillary. Expel a small drop of plasmid mix out of the tip to ensure no air is left behind in the tip.

26.1 The needle tip is fragile. Do not bump it on any surface or shift it during injections.

27 Use your non-dominant hand to stabilize one embryo between thumb and index finger. The embryos can be slightly manipulated within the amniotic pouch to better orient embryos for injection.

27.1 The ideal position for injection is for the embryo's head to be lightly pressed up against the inside of the uterine wall. This stabilizes the embryos within the sac and minimizes the distance needed for the injection step.

27.2 Do no over manipulate as it may rupture the amniotic sac or damage the placenta or the surrounding blood vessels.

28 Identify the 3rd ventricle. It will be slightly more posterior than the most dorsal point of the embryo.

29 Holding the microcapillary close to the needle tip, gently poke into the most caudal region of the 3rd ventricle.

29.1 The tip depth of the needle into the skull if very short as the ventricle is located close to the surface.

29.2 You may feel two parts to this injection - first the needle puncturing the uterus and then the needle entering the embryo's head. However, with a sharply beveled microcapillary you may not feel much resistance at all. If resistance is high during injection, the injection needle is too dull and should be replaced.

29.3 The Fast Green dye allows for visualization of the injection mix in the ventricle. If you do not see the dye immediately filling the ventricle, the injection angle or depth is likely incorrect. Do not make more than one attempt per embryo.

30 Slowly inject 1 µL of plasmid mix into the 3rd ventricle until you see the 4th ventricle filling. Then gently pull out the microcapillary. Repeat this process for all of the viable embryos. 5m

30.1 Filling the 4th ventricle through the 3rd ventricle (via the cerebral aqueduct) greatly improves survival over direct injection into the 4th ventricle.

30.2 Inexperienced surgeons should opt for alternating between injecting the plasmid mix and electroporating the embryo. If too much time passes between injection and electroporation, the plasmid mix may diffuse and become diluted, which will affect electroporation efficiency.

31 Wet the uterine horns and the electrodes with HBSS.

32 Arrange electroporator foot pedal so that you can easily find it with your foot for the next step.

33 Place the cathode tweezer electrodes on the lateral sides of the embryo's upper neck, just ventral and slightly caudal to the 4th ventricle that filled with Fast Green dye. Place the anode 5m

single stick electrode directly on the dorsal side of the 4th ventricle, right on top of the Fast Green dye.

34 Press the foot pedal and wait until the electroporation to end, signaled by a double beep at the end of the 4 pulses.

34.1 You should see the embryo contract slightly during the electroporation. After the electroporation there should be slight scarring on the uterus where the electrodes were.

35 Injecting and electroporating the embryos should take no more than 10-15 minutes. If these steps are taking longer than this, skip some of the embryos to minimize time spent. The amount of time the surgery takes can greatly impact embryo/pup survival.

#### Suture and complete post-surgical care.

20m

36 When all embryos have been electroporated, carefully place uterine horns back into the abdomen. Dry the mouse abdomen with a Kimwipe to facilitate easier suturing and to keep the mouse warm. 30s

37 Using Graefe forceps in your non-dominant hand and needle holder in your dominant hand, perform a running suture along the muscle incision. Be sure to double knot at both the start and the end of the suture. 2m

38 Perform a running suture along the skin incision, starting and ending with double knots. 2m

39 Turn off the isoflurane but continue to allow oxygen to flow through the nosecone. Remove tape from paws. When the dam begins to wake up, quickly move to prepared cage resting on the heating pad. 30s

39.1 It is important that the cage contain enrichment such as cotton, paper for nesting, and plastic hut. Environment enrichment is essential for proper embryo development and vital for postnatal survival. Other enrichment such as sunflower seeds and breeder chow can also improve postnatal survival.

40 Allow mouse to wake up in cage in warm environment. Within 5 minutes the dam should be moving around. 5m

41 Place cage back on rack to allow mouse to drink water and check back 10 minutes later to observe the mouse. 10m

42 Inspect the mouse daily for two days to inspect the wound and assess pain. Administer and record additional doses of Meloxicam, if indicated by pain assessment. Pay special attention to see if stitches are removed.

42.1 Because excessive stress to the mother can result in birthing complication, litter rejection and cannibalism, try to place cage in a room with less traffic. Disturb cage as little as possible until a day or two after birth.

43 Mothers will give birth on the 19th day of gestation, typically in the early morning hours.

#### Protocol references

Meyer-Dilhet, Géraldine, and Julien Courchet. “In Utero Cortical Electroporation of Plasmids in the Mouse Embryo.” *STAR protocols* vol. 1,1 100027. 19 Jun. 2020, doi:10.1016/j.xpro.2020.100027

#### Acknowledgements

We thank Dr. Adam Green, Dr. Siddhartha Mitra, and Salvador Guerra for helpful scientific discussions.
